# Reconciliation of theoretical and empirical brain criticality via network heterogeneity

**DOI:** 10.1101/2021.03.11.435016

**Authors:** Lei Gu, Ruqian Wu

## Abstract

Inspired by heterogeneity in biological neural networks, we explore a heterogeneous network consisting of receipt, transmission and computation layers. It reconciles the dilemma that the data analysis scheme for empirical records yields non-power laws when applied to microscopic simulation of critical neural dynamics. Detailed analysis shows that the reconciliation is due to synchronization effect of the feedforward connectivity. The network favours avalanches with denser activity in the first half of life, and the result is consistent with the experimental observation. This heterogeneous structure facilitates robust criticality against external stimuli, which implies the inappropriateness of interpreting the subcritcality signature as an indication of subcrtical dynamics. These results propose the network heterogeneity as an essential piece for understanding the brain criticality.

---

The power law statistics for spike avalanches in *in vitro* [1, 2] and *in vivo* [3–7] observations is promoting the brain criticality from a hypothesis [8, 9] to a reality of cortical states, which is supposed to bear optimal capacity for information processing [10–18]. While the size distribution *p*(*s*) ∝ *s*^−3*/*2^ and duration distribution *p*(*t*) ∝ *t*^−2^ accord with the critical branching process, understanding of the origin of these power laws is still vague, even on the appropriate definition of the avalanche. A recent critique [19] calls into question the definition based on summing activities above a threshold. It shows that some widely used models compatible with this definition are in the universal class of random walk rather than the branching process. More seriously, in the study of branching process and analysis of experimental data, two distinct definitions are usually adopted [see Fig. 1(a) and the caption for detail]. In theoretical simulation, the cause (another spike or external stimuli) for each spike is tracked, and avalanches can be defined with causality [20–23]. In experimental measurements, the capacity for this detailed tracking is unavailable. As a workaround, the avalanches are identified with the time-bin scheme [24, 25]. Fig. 1(a) illustrates that the causal and time-bin avalanches are different for the same time series of spikes. Ref. [26] highlighted the dilemma between the two definitions, and raised the inconsistency that the critical branching process may not yield the observed power laws.

**FIG. 1.**
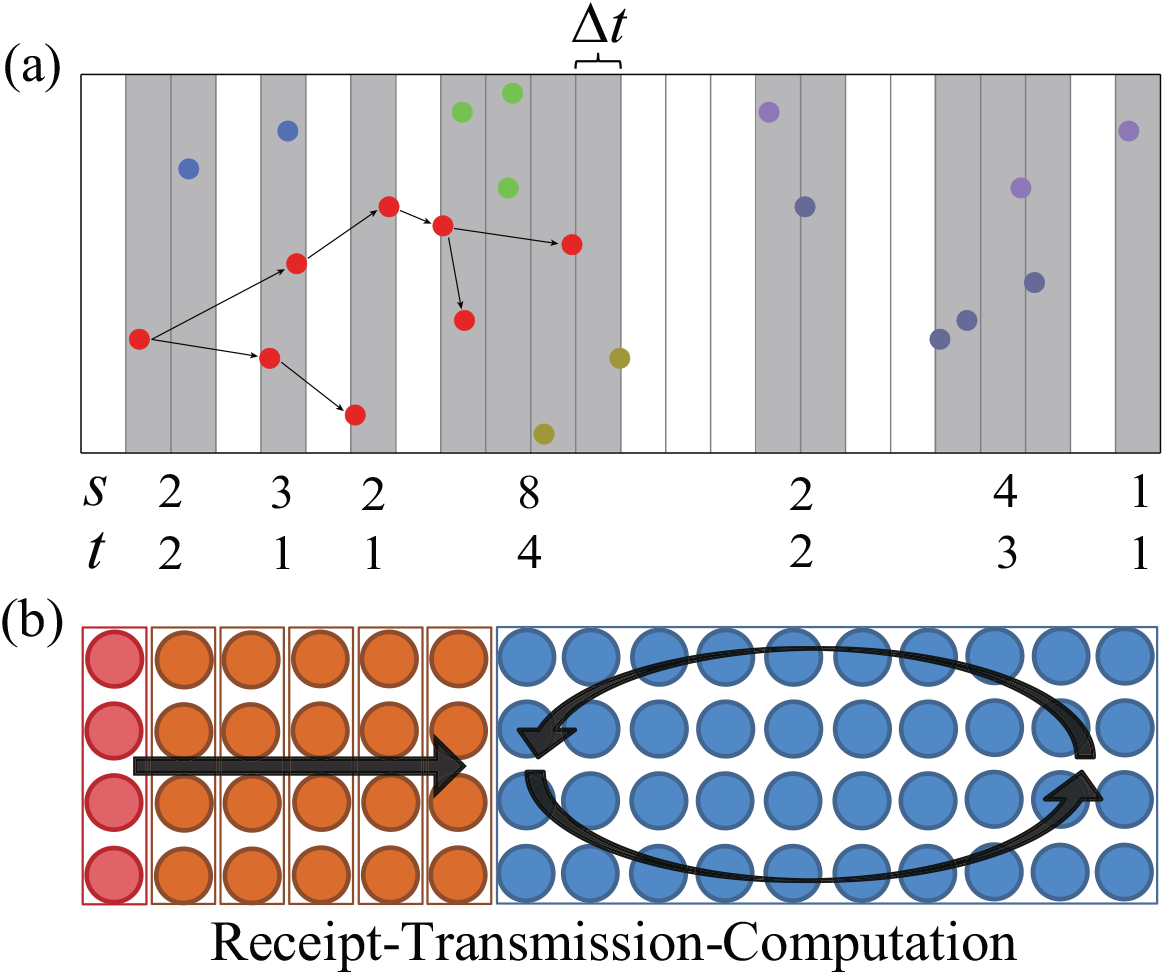
(a) In the time-bin definition of avalanches, an avalanche is defined as a cluster consisting a continuous sequence sandwiched between two empty bins (the shaded areas); The *s, t* numbers denote the sizes and durations. For the definition based on causality, the causal spike for a descendent spike is identified. Until an external stimulation brings in external cause, all descendants of an original spike (including itself) form an avalanche. Their size and duration are the number of spikes (dots in an identical color) and the corresponding time span. (b) The receipt-transmission-computation architecture consists of an input layer, feedforward layers for signal transmission, and a network of recurrently connected neurons modeling the cortical layers.

The gist of this work is the finding that the feedforward connectivity furnishes an effective synchronization mechanism. By building a heterogeneous network architecture made of receipt, transmission, and computation layers, the dilemma of avalanche definition can be reconciled, that is, the two definitions both produce correct power laws. Here, the computation part models the cortex, where the activities are recorded and the criticality can be found. The synchronized activities from the transmission part tend to induce avalanches in the computation part that have dense spiking rate in the early stage, which is consistent with experimental observations. Moreover, such a heterogeneous structure imparts a functional advantage in that the criticality is robust against varying external inputs. These results substantially extend the existing understanding based on homogeneous systems [4, 27–40].

While the network heterogeneity receives little attention in studies of the brain criticality, it plays a central role in the research of brain functionalities [41–44], as a featureless homogeneous network is incapable of mimicking meaningful tasks. For instance, the visual path way involving retina neurons, the lateral geniculate nucleus, and the visual cortex [45] constitutes a typical receipt-transmission-computation architecture (RTC) for signal processing. In terms of cortical layering, it is observed that certain layers are main targets of inputs, and neurons in some layers are more recurrently connected [46, 47]. The network structure throughout this work is a simple mimic of the RTC architecture as sketched in Fig. 1(b). Neurons in the first layer are recipients of external inputs. Ensuring are feedforward layers that transmit the signals to the computational region, which is made of randomly connected neurons (an Erd ő s-R ényi network). We use the leaky neuron model proposed in Refs. [20–22] as the building blocks, since it gives a rather practical description of neuronal dynamics and accommodates stochastic vesicle release, self-organized criticality [20, 21], and up-and-down cortical states [22, 48].

Fig. 2(a) demonstrates the reconciliation between the causality and time-bin definitions of avalanches, where both give the correct power laws. The most direct cause for the reconciliation could be temporal separation of the causal avalanches. Without the overlaps in time and with proper time bin width, the two definitions would lead to similar clustering of spikes. From the excerpt in Fig. 2(b), however, the avalanches (denoted in different colors) are highly overlapping. An alternative cause could be the criticism [49] for criticality that the power law statistic of time-bin avalanches is a general consequence of setting large scale networks in the Boltzmann’s molecular chaos regime [50]. According to this criticism, the criticality is not necessary for the power law statistics. However, we find that once presented, in this heterogeneous structure the criticality is a robust presence against external inputs.

**FIG. 2.**
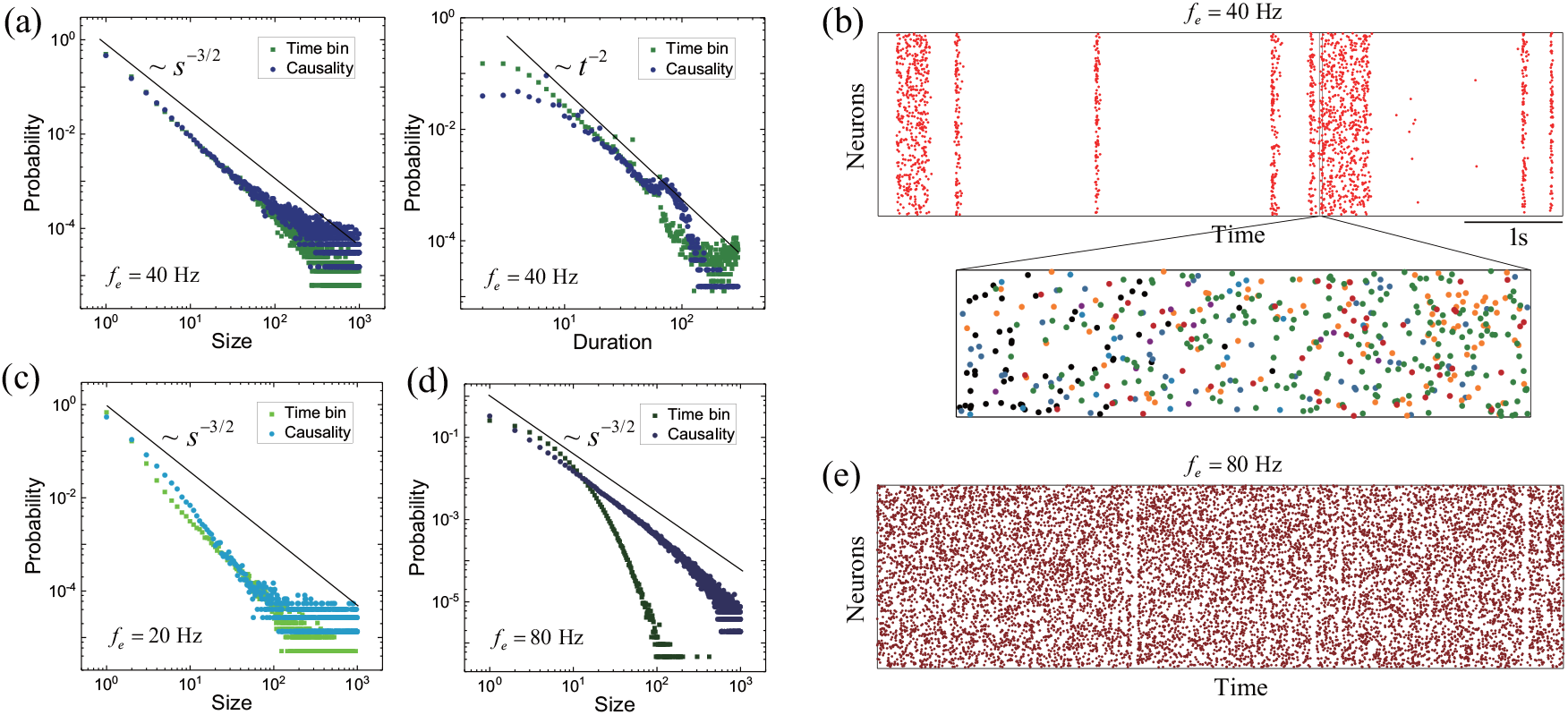
(a) The two definitions of avalanches both give the correct power laws; here *f*_*e*_ = 40 Hz. (b) Synchronization leads to bursting activities; The room in shows that the causal avalanches are temporally overlapping, where the black includes avalanches having *<* 10 spikes. (c) Lowering andincreasing frequency of the inputs do not obviously deviate the dynamics from the critical point. (d)High input frequency compromises the synchronization effect and results in the non-power law in (d).

Turning down the frequency of the input to *f*_*e*_ = 20 Hz, we have the size distributions in Fig 2(c). The branching rate *σ*_*c*_ = 0.99 (subscript *c* means it is based on causality; see the next paragraph) indicates that the dynamics is quite close to the critical point. For high frequency *f*_*e*_ = 80 Hz, the branching rate is *σ*_*c*_ = 0.96. The size distribution of the causal avalanches is the deep blue curve in Fig. 2(d), and it shows that the criticality is only slightly compromised. Comparison between the time-bin and causality curves suggests that the non-power law for the time-bin avalanches is due mainly to ineffective synchronization [cf. Fig. 2(e)], and due much less to the deviation from critical dynamics. When the frequency of input further increased, the branching dynamics can obviously deviating from the critical point [51]. Except for such extreme situations, Fig. 2(d) suggests that the observed subcriticality signature should not be interpreted as an indication of subcrtical dynamics.

To prevent frustration, we remark on the definitions of branching rate in the causality and time-bin schemes. For modeling of branching processes, the branching rate is defined as the average of *σ*_*i*_ = ∑_*j*_ *p*_*ij*_, where *p*_*ij*_ represents the probability of item *i* generating item *j* in a propagation step. The branching rate based on causality is defined as *σ*_*c*_= ∑_*i*_ *σ*_*i*_*/N* for a network of *N* neurons, with 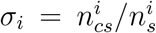, where 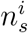 and 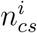 denote the number of spikes for neuron *i* and the number of spikes caused by these spikes. This definition is instantiation of the probabilistic definition. In the time-bin scheme, the branching rate is defined as *σ*_*t*_ = ∑_*n*_*σ*_*n*_*/*(*T* − 1) for a time series of *T* bins, where *σ*_*n*_ denotes ratio between the number of spikes in the (*n* + 1)th and the *n*th bin. As the causal spike and the caused spike are not necessarily in adjacent bins [cf. Fig. 1(a)], *σ*_*t*_ can be quite different from the standard definition, and thereby is an inaccurate characteristic for the branching dynamics. On the contrary, *σ*_*c*_ ≈1 is an indication of branching dynamics close to criticality. Since the criticism of Ref. [49] is based on the time-bin scheme, it does not compromise the validity of the power law statistics for the casual avalanches as an indication of critical dynamics.

To demonstrate the crucial role of the transmission layers, we try a structure where the receipt layer is directly connected to the computation part. As shown in Fig. 3(a), the size distribution of both the time-bin avalanche and causal avalanche obviously deviates from *s*^−3*/*2^. The corresponding branching rate *σ*_*c*_ = 0.65 also indicates subcritical dynamics. To confirm that the transmission layers indeed furnish a synchronization mechanism, we compute the variation of spiking rate for each layer, which characterizes the temporal concentration of activities. More straightforwardly, for each layer we also count how many spikes occurs within a ±10 ms around a certain spike. As shown in Fig 3(b), both quantities show monotonic increase when going deeper. These results suggest that the deeper a layer is, the neurons in that layer tend more to spike together. The computation part itself is an homogeneous network receiving inputs from the transmission part. In contrast to the opinion that inputs drive the dynamics away from the critical point [22, 24, 47, 52], comparison between Fig. 3(a) and the results in Fig. 2 suggests that the synchronized inputs can be an essential ingredient driving the dynamics to the critical point.

**FIG. 3.**
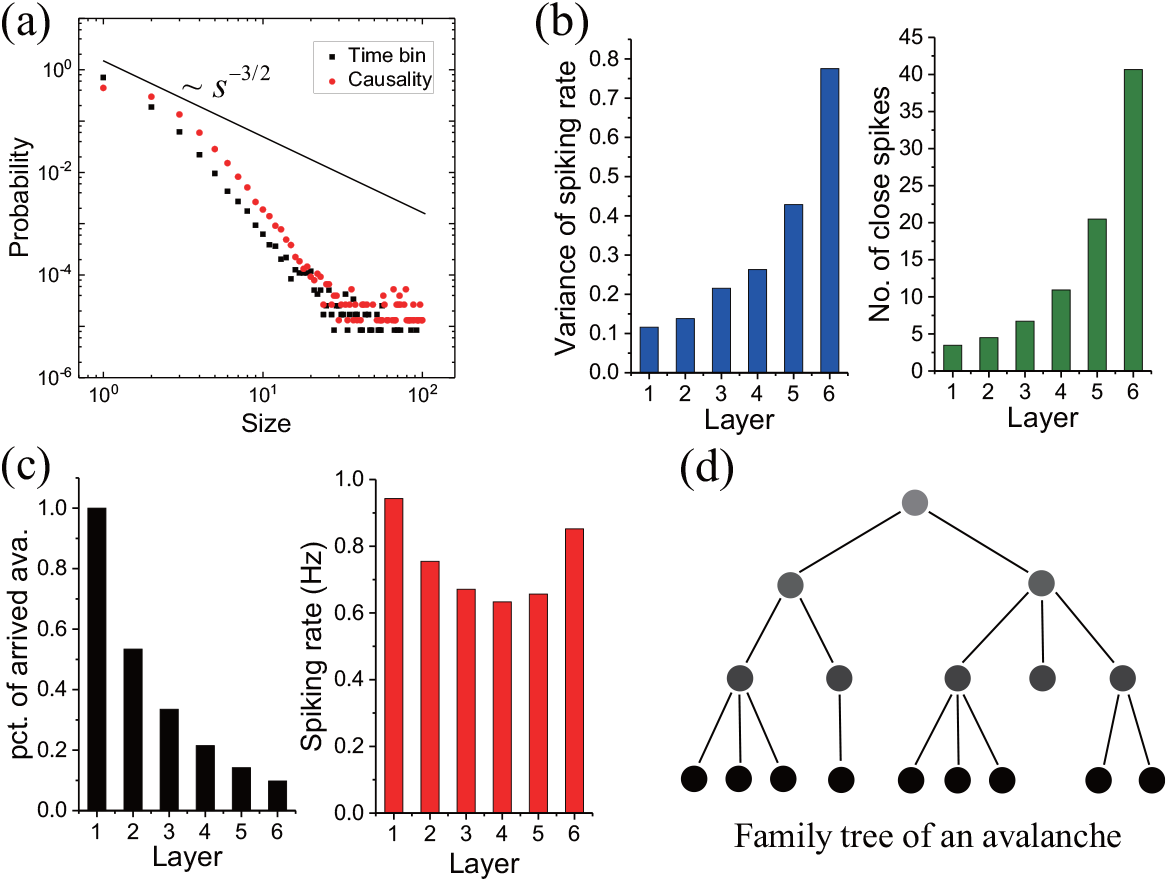
(a) Without the transmission part, the criticality is compromised and no correct power law can be rendered. (b) The variation of spiking rate and number of temporally close spikes monotonically increase with the layer number, indicating increasing synchronization of the activities. (c) Shape decrease of the proportion of arrived (causal) avalanches suggests that the transmission part is a low pass filter; together with less varied spiking rate, a family tree with increasing descendants can be inferred for the survived avalanches, as the schematic of (d).

The reason for the synchronization can be inferred from the proportion of arrived causal avalanches and the average firing rate at each layer [Fig. 3(c)]. The dramatically decreasing number of arrived avalanches implies that many avalanches die out in every propagation step. Together with the less varied firing rate, it can be deduced that the survived avalanches have (in average) more spikes in a layer when going deeper. In other words, in the transmission part a parent spike tends to have multiple offspring spikes. Because of the feedforward connectivity, descendant spikes in a layer belong to the same generation, as depicted in the family tree schematic of Fig. 3(d). In analogue to the phenomena that descendants of an ancestor that are in the same generation tend to have closer birth dates than randomly selected people, spikes belong to the same avalanche and in the same layer have closer spiking time than independent Poisson processes. Because the postsynaptic incentive diminishes quickly, a spike have high probability to produce offspring only in a short period of time. Accordingly, the synchronization effect is quite obvious.

Besides synchronization, another effect rendered by the feedforward connectivity is of functional importance. We find that a feedforward layer can function as an amplifier when firing rate of its input exceeds a threshold. Namely, the firing rate of a layer is larger than the preceding one, if the firing rate of this preceding layer is high enough. The inverse is also found to be true. As a result, controling the firing rate of the first layer, one can have the monotonically increasing or decreasing firing rate, as shown in the top and down panels of Fig. 4(a). We also identify this bifurcation in the analytical solution of the model as shown in Fig. 4(b). Experimentally, amplification of activities crossing a threshold has been observed in various brain areas [53]. A circuit of recurrent feedback based on a linear integrate-and-fire model was designed to account for this [54]. Here, using the more practical neuron model, we show that feedforwad connectivity is sufficient to give rise to the phenomenon. Its role in selective amplification of signals [55] is an interesting topic for further studies.

**FIG. 4.**
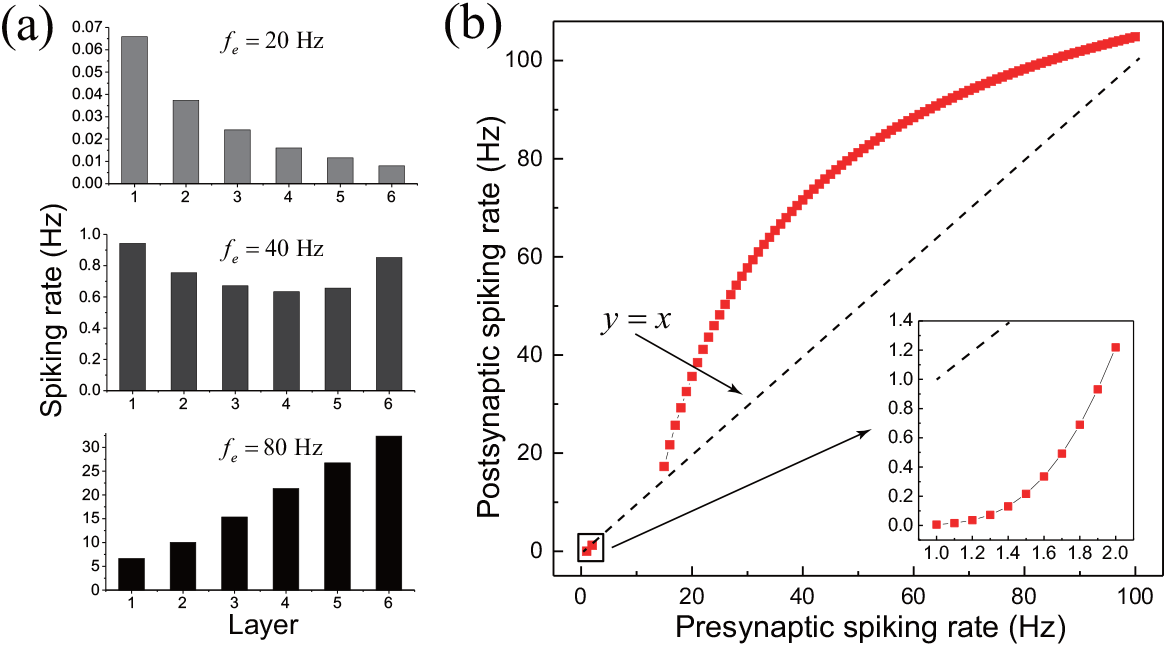
(a) Simulation results for inputs with three different frequencies. With variation of firing rate of the first layer, three patterns of the layer-wise spiking rate can be achieved in the transmission layers; the case of high frequency means that feedforwardly connected layers can be used as an signal amplifier. (b) the bifurcation is also present in the analytical solution; the post- and pre-synaptic spiking rates are the average firing rate of neurons in the layer being considered and the preceding layer, respectively.

In the synchronous irregular (SI) regime of the Brunel model [56], where the inhibitory neurons dominate over the excitatory neurons, the 3*/*2, 2 power laws can be produced with the time-bin analysis [49, 57]. In the RTC structure, the synchronization occurs in the transmission part, so it is external to dynamics of the computation part. For the inhibition-dominant Brunel model, the synchronization is due to inhibition effect of the inhibitory neurons, and thereby a built-in ingredient of the neural dynamics. We compare simulation results of these two models with the experimental data of Ref. [58] to demonstrate what observational signature the RTC structure brings about.

As shown in Fig. 5(b), both models generate power law statistics (time-bin) comparable to the empirical data. Fig. 5(a) suggests that the neural activities take a bursting pattern in all the datasets. To discern the distinction between the external and innate synchronization, we count the left and right inclined avalanches. Here,”left” and “right” denote whether the peak of activity for an avalanche locates in the first half of life or the second half of life. The postulate is simple that inhibition effect of the Brunel model relies on activity of the inhibitory neuron and in turn on the activity of the whole network, so the bursts tend to end with (relatively) dense spikes. In contrast, avalanches in the computation part of the RTC network is initialized by synchronized stimuli, which may induce dense activities in the early stage. We plot the proportion of the left inclined avalanches versus avalanche size in Fig 5(c). For the most part, the experimental data and the RCT result have proportions *>* 0.5, and vice versa for the inhibition-dominated network (noted as “IN”). We show in the supplementary that these trends are robust to variation of model parameters and subsampling of the RTC data yields results more consistent with the experiment data.

**FIG. 5.**
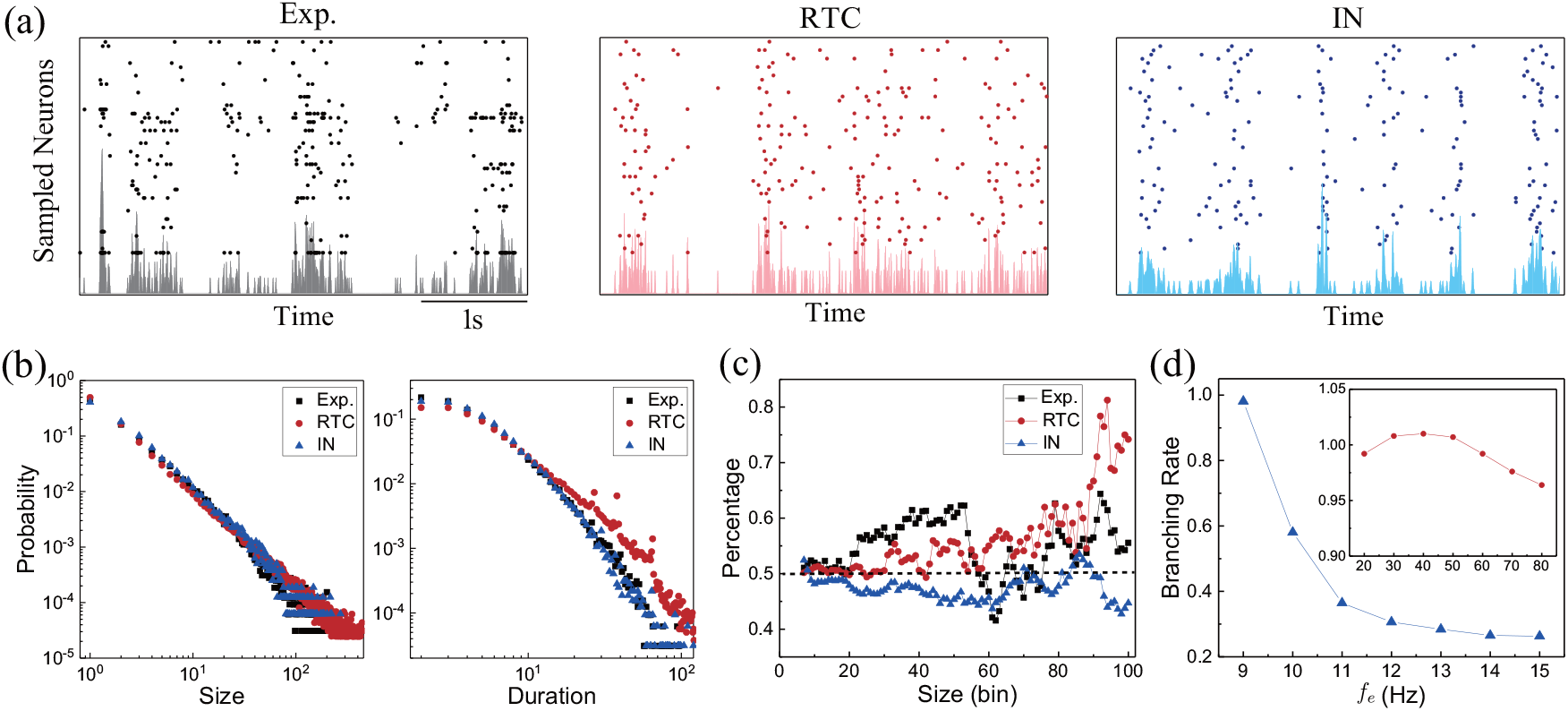
(a) Excerpts of experimental records from Ref. [58] and simulation results. (b) Both the models generate power law statistics comparable with the experimental data. (c) Proportion of the left inclined avalanche; The RTC network has proportion *>* 0.5 for most sizes, better consistent with experiment. (d) The branching rate of the Brunel model shapely decays when going away from the lower boundary of the SI regime, while branching rate of the RTC network is much insensitive to external stimuli.

With our parameter setting for the Brunel model, the threshold frequency is *f*_*thr*_ = 12.5 Hz (see supplementary for details), and the SI regime corresponds to 0.74 *f*_*ext*_*/f*_*thr*_ 1.1[56]. With the causal branching rate, we can peer into the underlining dynamics. From the branching rate for *f*_*e*_ = 9 ∼ 15 Hz in Fig. 5(d), there appears to be a phase transition near *f*_*e*_ = 9 Hz. Away from it, the decreasing branching rate indicates rapid approaching to subcritical regime. In contrast, the inset shows that the dynamics of the RTC network is much more robust against the variation of external stimuli. As the inhibitory neuron is a recognized component of the biological neural network, and rich repertoire of states of the Brunel model provides description to diverse activity patterns of neural networks, we aim not to repudiate them. Rather, due to its account for the favorable avalanche pattern, we propose the RTC structure as a necessary mechanism of synchronization. In addition, this comparative investigation raises a general caveat that the power law statistics is insufficient for characterizing the neural dynamics, since the firing pattern can be quite different.

In conclusion, by prompting the homogeneous network structure to the more biophysically practical heterogeneous structure consisting of receipt, transmission and computation layers, the dilemma between the time-bin and causality definition of avalanches can be reconciled. The synchronized stimuli to the computational part yield the favorable avalanche pattern consistent with the experimental records. The variability of spiking patterns under the same power laws implies that the power law statistics is insufficient and should be complimented by the firing pattern for better characterizing the neural dynamics. The heterogeneity also facilitates robustness of the criticality against variation of external stimuli, which call into question the conventional interpretation of the observed subcritical signature. Ineffective synchronization rather than subcritical dynamics should be the major cause for this signature. Most of all, the network heterogeneity should constitute an essential piece in our understanding of the criticality in brain.

We thank J. M. Beggs for explanation to the data processing procedure for experimental measurements, J. Hidalgo for advices on the numerical simulation, and Y. Senzai for support with the experimental data. This work was supported by the National Science Foundation through the University of California-Irvine Material Research Science and Engineering Center (grant no. DMR2011967). Codes for the implementation and analysis of the Millman model are available at https://github.com/Lei-Gu-901/Millman.

^∗^

## Supporting information

Supplemental results and discussions

