## Supplemental results and discussions for "Reconciliation of theoretical and empirical brain criticality via network heterogeneity"

### Supplementary information for "Reconciliation of theoretical and empirical brain criticality via network heterogeneity"

Lei Gu and Ruqian Wu*

*Department of Physics and Astronomy,*

*University of California, Irvine, California 92697, USA*

#### INTRODUCTION TO INVOLVED NEURON MODELS

##### The Leaky neuron of Millman *et al.*

In this model, a neuron receives currents from and sends currents to $n_c$ other neurons, respectively. For a neuron (in terms of presynaptic), the neurons being connected to (postsynaptic) are randomly selected. That is, the connections form an Erdős-Rényi network. Signals are transferred through vesicle release by $n_r$ release sites on each synapse. When a neuron spikes at time $t^k$, a vesicle release is launched with a certain probability, which injects a current $I_{in}^k(t) = w_{in}e^{-(t-t_k)/\tau_s}$ into the postsynaptic neurons. Every neuron also receives independent Poisson external currents $I_e$ with amplitude $w_e$, frequency $f_e$ and the decay time $\tau_s$. Synaptic depression is introduced in by time-dependent utility for each release site, which is set to $U = 0$ at the launching time and recovers exponentially toward 1 with characteristic time $t_R$, i.e., $U(t) = 1 - e^{-(t-t^k)/\tau_R}$. When a neuron spikes, the probability of launching a vesicle release is $p_rU(t^k)$, where $p_r < 1$ is a constant and $t^k$ the spiking time.

Putting these ingredients together, the membrane potential of the $i$th neuron obeys the equation

$$\dot{V}_i(t) = -\frac{V_i - V_r}{RC} + \sum_k \frac{I_e^{k_i}(t)}{C} + \sum_{k,s,j} \frac{1}{C}\Theta\left(p_rU_{js}(t^{k_j}) - \zeta_{js}\right) I_{in}^{k_j}, \tag{1}$$

where $R$ is the membrane resistance, $C$ the capacitance, $\Theta$ denotes the step function, and $\zeta_{js}$ is a random variable uniformly distributed on $[0, 1]$. Index $j$ enumerates the presynaptic neurons of this neuron, and $s$ indexes release sites on the synapses thereof. Upon reaching the threshold voltage $V_\theta$, the neuron spikes. After a short refractory period $\tau_{rp}$, the $V_i$ is reset to the resting potential $V_r$, and during this period dynamics of the neuron is turned off. In general, we used the parameter values in Refs. [22] and [26]: $n_c = 7.5$, $R = 2/3$ GΩ,

$C = 30$ pF, $V_r = -70$ meV, $\theta = -50$ meV, $w_{in} = 50$ pA, $w_e = 95$ pA, $n_r = 6$, $t_{rp} = 1$ ms, $t_s = 5$ ms, $t_R = 100$ ms, $p_r = 0.3$.

##### The Brunel model

Here we give an introduction to the model and provide parameters we use. More details can be found in Refs. [49, 56]. In the Brunel model, the voltage of neuron $i$ is governed by the equation

$$\tau \frac{\mathrm{d}v_i}{\mathrm{d}t} = -v_i + R\tau \sum_{j=1}^{n} J_{ij} \sum_{k} \delta(t - t_i^k - D), \tag{2}$$

where $\tau$ denotes time scale of voltage decaying due to charge leaky, $R$ is a resistance-like quantity, $J_{ij}$ represents the current sent from $j$ to $i$ when neuron $j$ spikes at $t_j^k$, and $D$ in the $\delta$ function means the current injection actually takes place after a delay. For excitatory neuron $j$, $J_{ij} > 0$ and vice versa for an inhibitory neuron.

The network setting for the results in main text Fig.5 is as follows. The network consists of 5000 neurons, 1000 (20%) of which are inhibitory. These neurons are connected to 4000 external neurons, which are all excitatory and receives independent Poison current injection. A neuron has probability of $\varepsilon = 0.1$ to connects to a certain neuron (self-connection is excluded). The reset voltage is 10 meV. The threshold voltage for firing is 20 meV and $RJ_{ij} = 0.2$ meV for the excitatory neurons. Under these settings, the threshold input frequency is $f_{thr} = 12.5$ Hz. The synchronous irregular (SI) regime requires dominance of the inhibitory neurons. As they only make up 1/4 of the network neurons, their voltage injection $RJ_{ij}$ should be scaled by a factor of $g < -4$. With $D = 1.5$ ms, the frequency of the external stimuli corresponding to the SI regime is in the range $0.74 \lesssim f_{ext}/f_{thr} \lesssim 1.1$ [56].

##### The model of Shew *et al.*

Since this model and the alike are bases of many modelings and arguments for the functional optimality. We brief its form and connection with the above models. In this model, the probability of a neuronal firing is determined as

$$p_i(t+1) = \Omega_i(t) e_i(t) + \sum_j W_{ij}(t) s_j(t) \tag{3}$$

where $W_{ij}$ is the connection weight from neuron $j$ to neuron $i$, and $\Omega_i$ is the weight representing strength of external stimuli. $e_i$ and $s_j$ are indicators of the external stimuli and firing of neurons. $e_i(t) = 1$ if an external stimuli is performed upon neuron $i$ at time step $t$, otherwise $e_i(t) = 0$. It is similar for $s_j(t)$, which denotes whether neuron $j$ spikes or not at time step $t$. When $p_i > 1.0$, neuron $i$ spikes with probability of 1. When $0 < p_i < 1$, it spikes with probability of $p_i$. When $p_i < 0$, the neuron does not spike.

It is easy to see that when $\sum_j W_{ij} \approx 1$ and $\Omega_i \ll 1$ (weak external stimuli), a stationary firing rate of the neurons can be achieved. It means that the branching rate for every time step is $\sigma_s \approx 1$. We did not carry out comparative study with this type of models, since it is mostly a general modeling of branching processes rather than micro simulation of neural dynamics. Because caused spikes are at the time step following the causal one, causality is implicitly implemented in this model. There is no problem to calculate the branching rate as time average over $\sigma_n$, with $\sigma_n$ being the ratio between number of spikes in the $(n+1)$th and $n$th time step.

#### SUPPLEMENTARY RESULTS AND TECHNICAL DETAILS

##### More on the definition of causal avalanches

Since a neuron receives external stimuli and postsynaptic incentives of multiple neurons, accumulation of the membrane voltage involves all these stimuli, and the causal stimulus is identified as the one that makes the voltage finally cross the threshold. If it is an external stimuli, a new avalanche is initiated; If it is an postsynaptic current injection, the parent spike can be identified accordingly. For the computation part of the RTC network, we take the inputs from the last transmission layer as external stimuli. In other words, as long as neurons in this layer are identified as the causal neuron, a new avalanche is initiated.

##### Details of the RTC network

There is 1 receipt layer, 5 transmission layers, and 10 computations layers. Every layer has 81 neurons. Connections in the first 6 layers are fully feedforward. Namely, for every neuron (on average) 7.5 neuron in the following layer are randomly selected as targets of the postsynaptic connections. To render the computation part a homogeneous network (viewed along), 7.5 target neurons are random selected from all of the 10 computation layers. Here we stress that this setting is different from our prior work [51], where the connection with the last transmission layer is limited to the first computation layer. As shown there, doing this can further facilitate the robustness against external stimuli. With the current setting, the computation part itself is a homogeneous network.

Another difference should also be stressed. With parameter setting in the introductory section of the Millman model, the network is slightly supercritical (in the sense of independent Poisson stimuli), and weak inputs can lead to sustainable activities. As a result, the synchronization effect can not show up. To make the synchronization essential for sustainable activities in the cortical region and have the result presented in the main text, it is necessary to slightly tune the cortical region toward the subcritical regime. To this end, in the computation part we downscale $n_c$ by a factor of 0.9, that is $n_c = 6.75$ in this region. With such setting, synchronization of inputs (for the computation part) is an essential dynamical ingredient for reaching criticality. This point implies that the existing studies based on homogeneous network and Poisson stimuli miss an important aspect. In a more integrated and complete theoretical consideration of the brain criticality, the degree of synchronization of external stimuli should be taken explicitly as a model parameter.

**Notes on width of the time bin**

The width of the time bin is usually determined as the average inter-spike interval of the time series. However, because there are long blank areas between burst in the cases of low frequency inputs, the interval that well reflect the neural activity should be much smaller than the interval of the whole time series. In this case, Ref. [1] suggests to determine a time window for each bursts according to the cross-correlation. To facilitate cross-frequency comparison, and because we have knowledge of the neuronal dynamics, for the Millman model we use $\Delta t = 100/810 = 0.123$ ms when the interval is larger than this value. Here 810 is the number of neurons in the computation part, and 100 ms is the recovery time of synaptic utility ($t_R$). We take this value as an estimation for the average interval after exclude the long inactive periods.

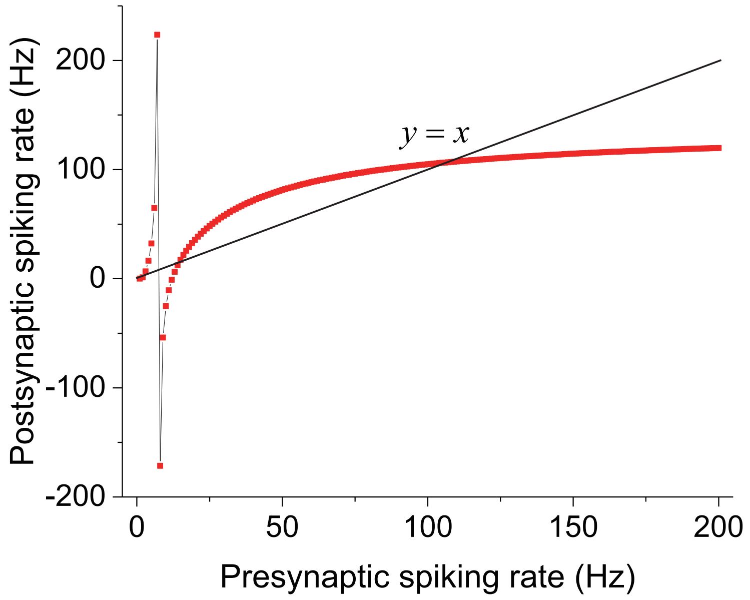


FIG. 1. Complete result of main text Fig.4. Because of simplification and impractical assumption in obtaining the Fokker-Planck equation, negative value and quantitative difference with the simulation result exist.

##### Complement results for main text Fig.4

The solution of the Fokker-Planck equation is given in the Appendix of our prior work [51]. In Fig. 1 we complement the result in main text Fig.4. In the transition regime, negative frequencies can exist, and this is nonrealistic. When the presynaptic rate exceeds the crossing rate( i.e., $> 106$ Hz), it would rapidly decrease to $\approx 106$ Hz. This explains why the firing rate shapely declines with the propagation of neural activities [51]. There the simulation result gives a crossing spiking rate $\approx 70$ Hz. The major reason for the negative frequencies and the quantitative difference could the independence assumption in deriving the Fokker-Planck equation. Namely, one should assume that the activity of each neuron is independent Poisson process. However, as the spikes are caused (mainly) by postsynaptic incentives, the activities of neurons are highly correlated. The rescaling procedure for postsynaptic current injection proposed in Ref. [48] can lower the crossing spiking rate as shown in [51]. Here we do not implement the procedure, since it introduces more nonlinearity and irregularity to the spiking rate.

**Complement results for main text Fig.5**

The experimental data of main text Fig.5 is from spiking records of mouse YMV08 of Ref. [58], which contains activity records of 101 probe channels. The data processing procedure of Ref. [6] is performed upon the data. The power laws are derived from the 50 10s-blocks with the largest coefficient of variance, since they yield exponents close to the critical ones (i.e., 3/2 and 2) [6]. The processing of dataset for 3 other mouses also follows this procedure. The width of the time bin is determined as average inter-spike interval, which is 2.7 ms for this mouse. Since the experimental dataset has 101 probe channels. The raster plots for the Millman and Brunel model are subsampling of about 100 neurons. We also properly rescale the plots to render similar looking with the experimental one, and the time-bar for the experimental data does not necessarily apply to the simulation results.

As mentioned in the section of Brunel model, the current injection by firing of inhibitory neurons should be over four times larger than that of excitatory neurons to reach the SI regime. The power laws for the Brunel is derived from simulation of $g = -4.1$ and $f_e = 12$ Hz. We choose this $g$ value to make the inhibitory neurons dominate but not overwhelming. The frequency of external stimuli $f_e = 12$ is away from the lower boundary (cf. Fig.2 of Ref. [56]) of the SI regime so that the activity is not too sparse. Since $g = -4.1$ is too close to the phase boundary, we find that the variation of branching rate $\sigma_c$ for the Brunel model is too large in this case. To have stable results with small variation, we set $g = -5.1$ in obtaining the results in main text Fig.5(d).

In the following, we show the robustness of the trends in main text Fig5(c). First, we should note that the value for every size in Fig.5(c) is an average over 9 sizes (the size itself being the center and the $4+4$ nearby sizes). The data without the average is given in Fig.2, where the fluctuation is quite strong. The average smooths the fluctuations and makes the trends clearer.

The width of the time bins for plots in main text Fig.5(c) are not the average inter-spike interval ($\Delta t$), but twice of it ($2\Delta t$). We show in Fig. 3 that the trends are not qualitatively changed by the variation of bin widths ($\Delta t, 2\Delta t, 3\Delta t$). The preference for the left-incline avalanches is made clearer with increasing width, especially for the experimental data. Basically, since the synchronized inputs from the transmission part induce bursts containing multiple avalanches, what we actually needed to discern is the left-inclined and

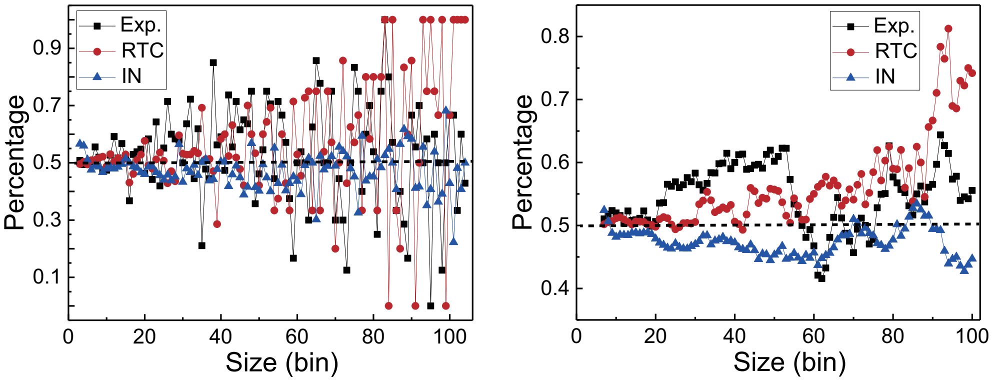


FIG. 2. Left: the data for main text Fig.5(c) without average with nearby sizes; the fluctuation is strong and it is difficult to draw a clear trend. Right: repetition of main text Fig.5(c)

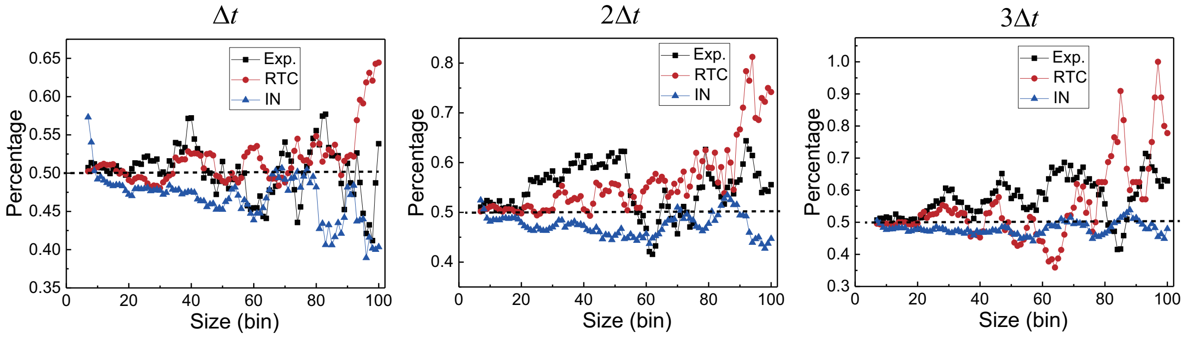


FIG. 3. Proportion of the left-inclined avalanches with varying width of the time bin. No qualitative change occurs. For the experimental data, larger bin width makes the preference of the left-inclined avalanches clearer

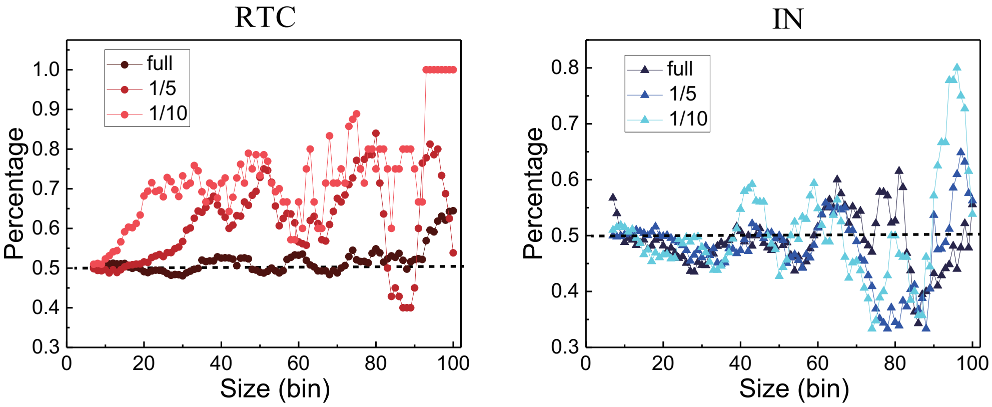


FIG. 4. Proportion of the left-inclined avalanches under subsampling. The RTC results become more similar with experimental data. The trends of the Brunel model's results do not change. Note that sparse subsmapling can cause large fluctuation.

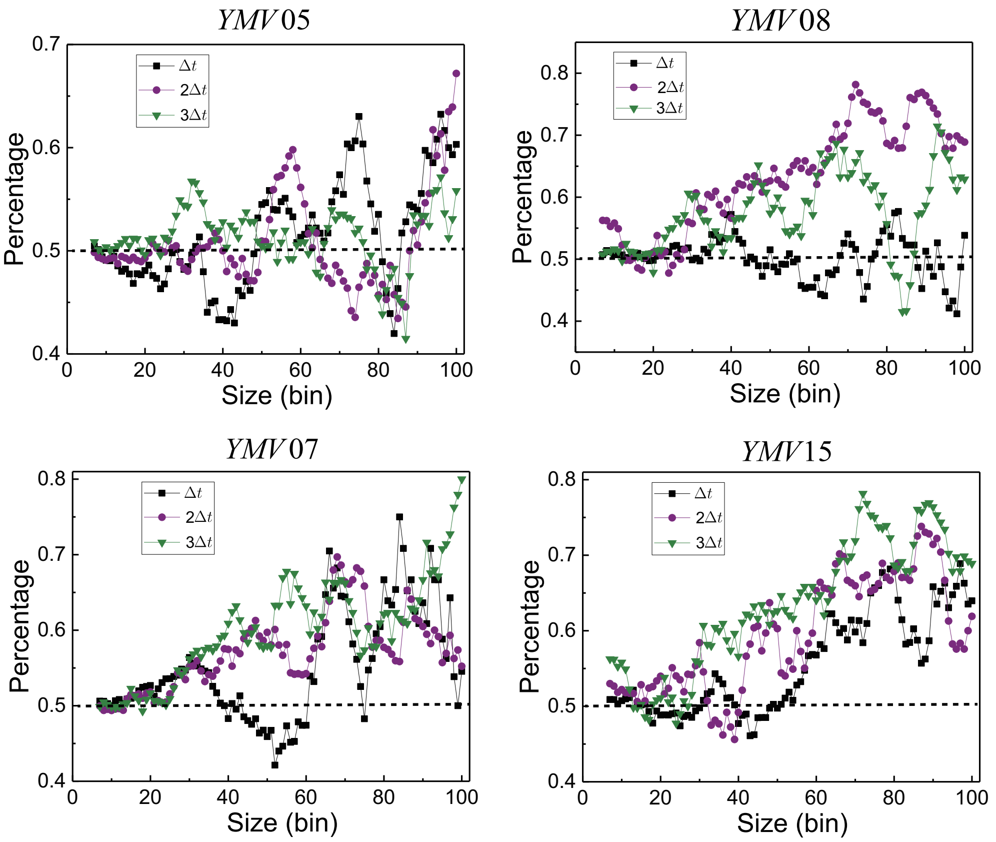


| Mouse | 05 | 07 | 08 | 15 |
|---|---|---|---|---|
| No. of probe channels | 74 | 102 | 101 | 81 |
| Average interval (ms) | 1.3 | 2.0 | 2.7 | 2.2 |

FIG. 5. Proportion of the left-inclined avalanches of 3 other mouses, together with that of YMV08. The trend of having proportions $> 50\%$ is clear, especially with enlarged bin width. The YMV05 might be an outlier that has neural activity twice stronger than other mouses.

right-inclined bursts. Increasing the bin width may lead to better identification of bursts and hence the trends are make clearer.

Since the subsampling issue is an inherent aspect of experimental records. In Fig. 4 we illustrate the effect of subsampling. With subsampling, the RTC results are even more similar with the experimental results. Meanwhile, these is no clear change of the Brunel model's results. Because of reduced records, the subsampling amplifies fluctuations, especially for large avalanches, which are rare in itself.

To confirm that left-inclined avalanches are favorable accross samples, we analysis experi-

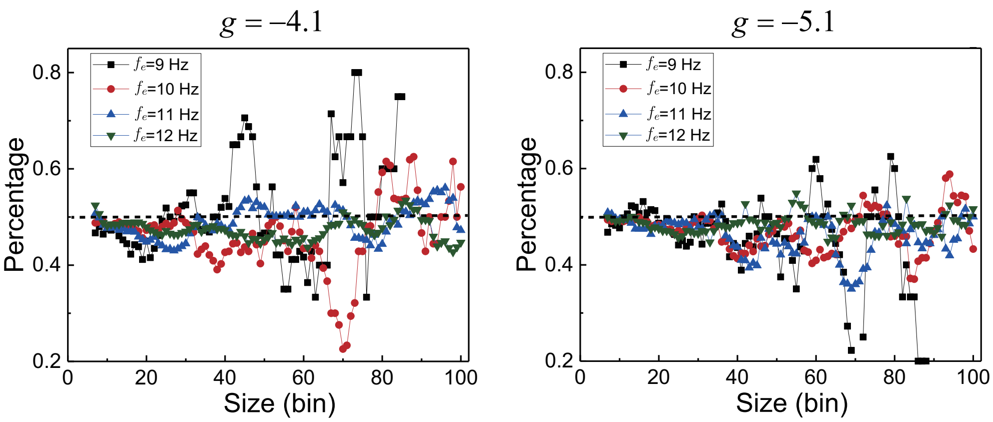


| $f_e$(Hz) | 9 | 10 | 11 | 12 |
|---|---|---|---|---|
| Spiking rate (Hz) | 4.81E-5 | 0.0043 | 0.012 | 0.019 |

| $f_e$(Hz) | 9 | 10 | 11 | 12 |
|---|---|---|---|---|
| Spiking rate (Hz) | 4.23E-5 | 0.0017 | 0.0051 | 0.0079 |

FIG. 6. Proportion of the left-inclined avalanches of the Brunel model with varying $g$ and $f_e$. The left-inclined avalanches are not favorable. Neural activity for $f_e$ near the lower boundary of the SI regime is very low.

mental data of 3 other mouses and show the results in Fig. 5. Our choices were depended on size of the datasets and not biased by our expectation. Namely, we selected 4 datasets whose number of probe channels are (relatively) high, and the time series are long. In general, the proportion of left-inclined avalanches are favorable. This trend becomes quite clear when the time bin width is augmented. In the table, we list the number of probe channels and mean spike interval. One see that mouse YMV05 has most sparse sampling and shortest spike interval. An estimation shows that its spiking rate is twice (or higher) of those of the others. This high activity may be the reason why there is no clear preference for YMV05, as the synchronization effect could be compromised or concealed. This mouse might be an outlier or there might be some subtle difference in the experimental set up.

To confirm that lower ratio of left-inclined avalanche is a general tread in the SI regime of the Brunel model. We plot the proportion for $f_e = 9, 10, 11, 12$ Hz and $g = -4.1, -5.1$ in Fig. 6. For the most part, the proportion $< 50\%$. The fluctuations for large sizes and low frequency $f_e$ is high, since low frequency of external inputs means sparse activity and large size avalanches are (relatively) rare. As shown in main text Fig.5(d), the dynamics is critical near the lower boundary ($f_e$) of the SI regime. In the table, we list the neuronal spiking rate for the parameter settings. The neural activity for low frequency inputs are quite low near the boundary ($f_e = 9.0$). This unreasonable sparse activity is another reason why this regime can hardly account for the brain criticality. In other word, the criticality of Brunel model in this regime requires very low activity (maybe even no activity at all) of the network.

---

*

[24] V. Priesemann, M. Wibral, M. Valderrama, R. Pröpper, M. Le Van Quyen, T. Geisel, J. Tri-

esch, D. Nikolić, and M. H. J. Munk, Spike avalanches in vivo suggest a driven, slightly subcritical brain state, Frontiers in Systems Neuroscience **8**, 108 (2014).

[37] S.-J. Wang and C. Zhou, Hierarchical modular structure enhances the robustness of self-

organized criticality in neural networks, New Journal of Physics **14**, 023005 (2012).

[50] L. Boltzmann, *Lectures on Gas Theory* (University of California Press, Berkeley, Los Angeles,

Chalifornia, 1964).
